# Reliable extraction of clinical-trial evidence with provenance, verification, and targeted review

**DOI:** 10.64898/2026.10.01.756091

**Authors:** Gaurav Najpande, Prasham Titiya, Muhammad Ali Khan, Kaneez Zahra Rubab Khakwani, Mohamad Bassam Sonbol, Irbaz Bin Riaz, Vivek Gupta

**Affiliations:** Arizona State University, Tempe, Arizona, USA; Mayo Clinic, Phoenix, Arizona, USA

**Keywords:** clinical evidence synthesis, information extraction, provenance, retrieval-augmented generation, human–AI collaboration, systematic review

## Abstract

Reliable extraction of trial characteristics and outcomes is essential for systematic reviews, yet incorrect values can alter evidence synthesis and are difficult to detect without clear source evidence. We evaluate EviSearch, a clinical evidence-curation system that links extracted values to source passages, independently verifies cited evidence, and prioritizes higher-risk cells for review. Using ten trial publications, a 133-column schema, reference annotations, and consistent scoring across model families, we assess answer quality, inter-reading agreement, evidence reproduction, targeted review, and evidence modality. In the Qwen evaluation, EviSearch achieved 91.70% accuracy, cited 100.0% of admitted values, and reproduced 98.9% independently. Reviewing 15.6% of cells increased accuracy to 95.22%.

## 1 Introduction

Systematic reviews and clinical practice guidelines depend on accurate information extracted from clinical-trial publications. Review teams must identify trial design, treatments, participant characteristics, outcomes, and harms across heterogeneous reports and changing terminology. This work is slow and error-prone, and its burden recurs as living reviews are updated [1–4].

Computational tools can reduce this burden, but clinical use requires more than a high average score. A treatment effect, eligibility criterion, or adverse-event count must be traceable to the publication and interpreted in the correct population, arm, timepoint, and unit. Unsupported values are costly to check, while remote processing can create governance concerns [5, 6]. Review definitions must also remain consistent as ambiguous reports are resolved.

We therefore evaluate clinical reliability directly: whether the evidence behind each value is visible, whether an independent check can reveal an unsupported reading, and whether expert review can be directed to higher-risk cells. Agreement and evidence reproduction are treated as risk signals, not guarantees of correctness or substitutes for clinical adjudication.

EviSearch [7] extracts structured trial evidence with source attribution and independent checking. Each value is linked to a page, quotation, and evidence modality, while cells with risk signals can be sent to a reviewer. Here we ask whether these safeguards remain useful across computational models, holding the curation task and scoring procedure fixed.

We test how extraction performance changes across model families, whether disagreement and failed evidence reproduction identify higher-risk cells, and whether source evidence can be independently reproduced across evidence modalities.

Recent studies support supervised computational assistance rather than replacement of clinical or methodological reviewers [8–11]. The remaining clinical gap is whether reviewers can verify extracted evidence and prioritize attention effectively [12–14].

Document-search and multimodal methods can locate evidence in full publications [15–17]. EviSearch requires the selected passage to support the precise clinical question and retains rendered pages for tables and figures that may be lost in extracted text.

Attribution and independent verification improve the safety of automated evidence extraction [18, 19]. EviSearch uses verification to inform adjudication and review rather than as an agreement gate, because failure to locate a value may reflect an incomplete reading rather than absence of evidence.

Human–AI review methods use uncertainty or disagreement to allocate expert attention [20, 21]. EviSearch applies this principle to complementary readings and stores versioned definitions and curation notes so review decisions remain consistent across evidence updates [22–25].

## 2 Results

### 2.1 Controlled model-family evaluation

The cross-model experiment changes the underlying language model while freezing the benchmark PDFs, schema version, knowledge snapshot, retrieval model, top-*k*, prompts, tools, reconciliation logic, verifier logic, and scoring pipeline. Dense retrieval uses Qwen3-Embedding-8B [26] with top-*k* = 5.

We evaluate four language-model families: Qwen3.6-27B [27], Mistral-Small-3.2-24B-Instruct-2506, Gemma-4-31B-it[28], and Llama-4-Scout-17B-16E [29]. For readability, these conditions are referred to throughout the manuscript as Qwen, Mistral, Gemma, and Llama, respectively. Debug and smoke runs are excluded from benchmark repetitions.

### 2.2 Established reference benchmark

The established Qwen benchmark provides the reference point for the journal evaluation. The single-pass baseline reached 89.01% accuracy, Agent A reached 89.71%, and Agent B reached 92.27%. Oracle selection of the better agent for each cell reached 93.21%. Full EviSearch reached 91.70% accuracy before simulated review, cited 100.0% of admitted values, and achieved 98.9% verifier reproduction. Simulated targeted review of 15.6% of cells increased accuracy to 95.22%.

Agent B was more accurate than EviSearch before simulated review in this benchmark. EviSearch, however, retained source attribution, independent reproduction information, and a review signal for the extracted clinical evidence. The following experiments examine whether these properties remain informative when the underlying model changes.

### 2.3 Cross-model generalization

Table 1 compares the single-pass baseline, the two extraction agents, and the final EviSearch output across four model families. EviSearch had higher point-estimate accuracy than the single-pass baseline for Qwen, Mistral, and Llama, while Gemma had lower accuracy. Qwen increased from 87.63% to 91.26%, Mistral from 79.17% to 81.52%, and Llama from 76.65% to 77.61%. Gemma decreased from 88.63% to 83.70%. The paired difference between EviSearch and B1 was 3.63 percentage points for Qwen with a 95% CI of 2.88 to 4.49, 2.35 points for Mistral with a 95% CI of *−*1.54 to 6.35, *−*4.93 points for Gemma with a 95% CI of *−*8.53 to *−*1.64, and 0.96 points for Llama with a 95% CI of *−*1.37 to 3.12. The interval was entirely above zero for Qwen and entirely below zero for Gemma, while the intervals for Mistral and Llama included zero. This shows that the effect of the EviSearch workflow relative to single-pass extraction differed across model families.

**Table 1.** Cross-model performance of the EviSearch pipeline across ten clinical-trial publications. Accuracy values are percentages. Confidence intervals are estimated using publication-clustered bootstrap sampling.

| Model | B1<br>Accuracy<sup>1</sup> | Agent A<sup>2</sup> | Agent B<sup>3</sup> | EviSearch<sup>4</sup> | E-B1<br>pp (95% CI)<sup>5</sup> | Papers<br>E > B1<sup>6</sup> |
| --- | --- | --- | --- | --- | --- | --- |
| Qwen | 87.63 | 89.14 | 91.05 | 91.26 | 3.63 (2.88, 4.49) | 10 |
| Mistral | 79.17 | 84.83 | 69.66 | 81.52 | 2.35 (-1.54, 6.35) | 6 |
| Gemma | 88.63 | 88.55 | 73.76 | 83.70 | -4.93 (-8.53, -1.64) | 3 |
| Llama | 76.65 | 82.69 | 49.77 | 77.61 | 0.96 (-1.37, 3.12) | 7 |

<sup>1</sup> B1 Accuracy is the accuracy of the single-pass baseline extraction, calculated as the mean of correctness and completeness across 1,330 cells.
<sup>2</sup> Agent A is the document-reading extraction agent, which extracts directly from the publication.
<sup>3</sup> Agent B is the search and retrieval extraction agent.
<sup>4</sup> EviSearch is the final reconciled output after comparison of Agent A and Agent B.
<sup>5</sup> E-B1 is EviSearch accuracy minus B1 accuracy in percentage points. The 95% confidence intervals were estimated using 10,000 publication-clustered bootstrap replicates.
<sup>6</sup> Papers E > B1 is the number of the ten clinical-trial publications for which EviSearch achieved higher paper-level accuracy than B1.

The paper-level results followed the same pattern. EviSearch exceeded the baseline on all ten Qwen papers, six Mistral papers, three Gemma papers, and seven Llama papers. The review queue captured 38.93% of EviSearch errors for Qwen, 72.05% for Mistral, 79.34% for Gemma, and 81.12% for Llama. These differences show that extraction performance and the concentration of errors requiring review vary across model families.

B1 denotes the single-pass baseline. Agent A extracts directly from the clinicaltrial publication, while Agent B uses document search and retrieval. EviSearch is the final reconciled output.

Figure 2 shows the uncertainty around the paired differences. The Qwen result was consistently above the single-pass baseline across the bootstrap interval, while the Gemma result was consistently below it. The wider intervals for Mistral and Llama included zero. The results show that the contribution of reconciliation to extraction accuracy was not the same across the tested model families.

**Fig. 1.**
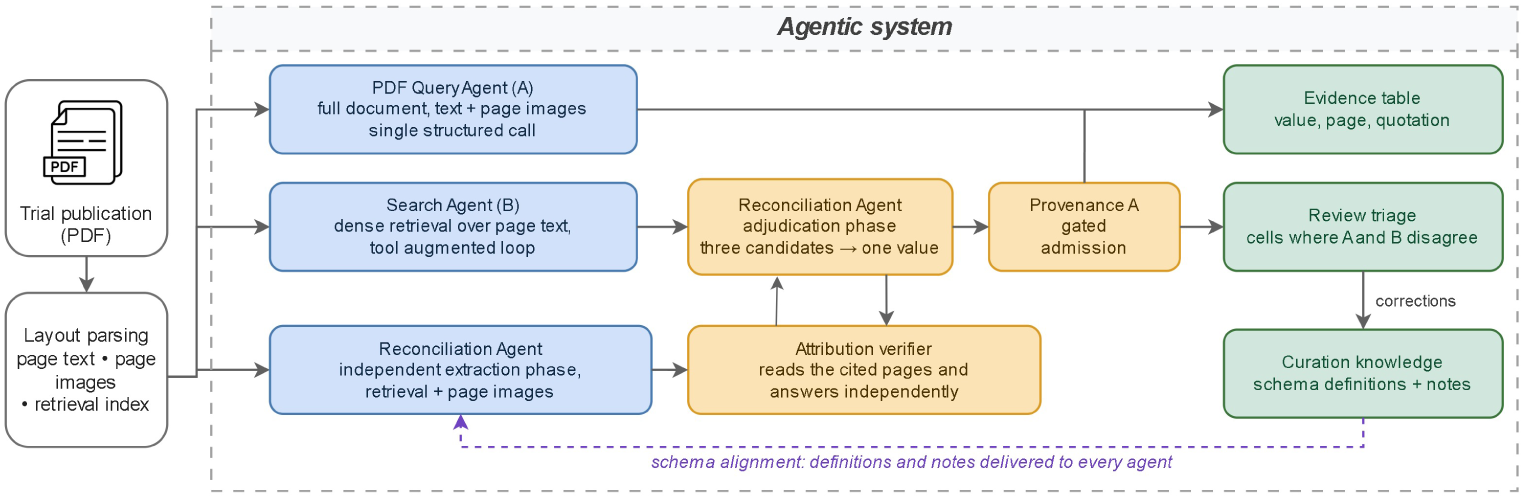
Architecture of EviSearch. Complementary extraction agents produce independent candidates; the Reconciliation Agent reads independently before adjudication; an attribution verifier checks the cited pages; and provenance-gated admission preserves the evidence and routes disagreement or weak corroboration to review.

**Fig. 2.**
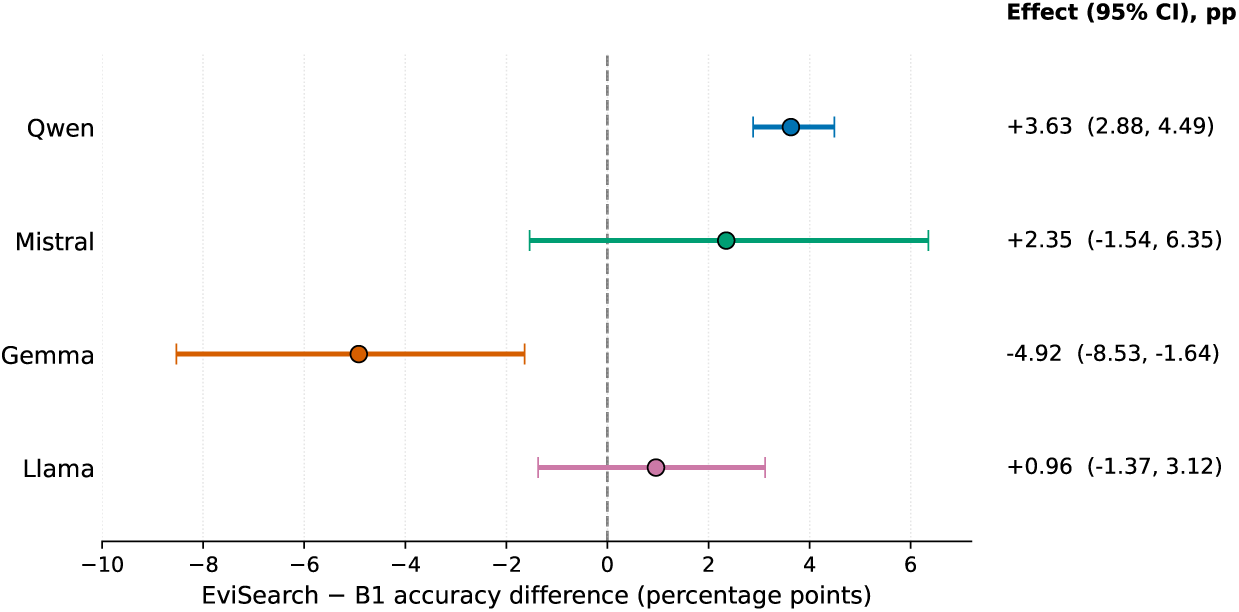
Paired difference in accuracy between EviSearch and the single-pass baseline across model families. Points show the EviSearch minus B1 difference in percentage points and horizontal bars show the 95% confidence intervals estimated using 10,000 publication-clustered bootstrap replicates. The dashed vertical line indicates no difference between EviSearch and B1.

### 2.4 Correctness and completeness

Table 2 separates accuracy into correctness and completeness. For Qwen, Mistral, and Llama, EviSearch improved both measures relative to the single-pass baseline. Qwen correctness increased from 88.05% to 91.80%, while completeness increased from 87.22% to 90.71%. Mistral correctness increased from 80.41% to 81.84%, and completeness increased from 77.93% to 81.20%. Llama correctness increased from 77.59% to 78.31%, while completeness increased from 75.71% to 76.92%.

**Table 2.** Performance decomposition across model families and extraction systems. All values are percentages averaged over 1,330 cells.

| Model | System | Accuracy (%) | Correctness (%) | Completeness (%) |
| --- | --- | --- | --- | --- |
| Qwen | B1 | 87.63 | 88.05 | 87.22 |
|  | Agent A | 89.14 | 89.62 | 88.65 |
|  | Agent B | 91.05 | 91.65 | 90.45 |
|  | EviSearch | 91.26 | 91.80 | 90.71 |
| Mistral | B1 | 79.17 | 80.41 | 77.93 |
|  | Agent A | 84.83 | 85.71 | 83.95 |
|  | Agent B | 69.66 | 70.19 | 69.14 |
|  | EviSearch | 81.52 | 81.84 | 81.20 |
| Gemma | B1 | 88.63 | 89.10 | 88.16 |
|  | Agent A | 88.55 | 88.83 | 88.27 |
|  | Agent B | 73.76 | 74.02 | 73.50 |
|  | EviSearch | 83.70 | 83.76 | 83.65 |
| Llama | B1 | 76.65 | 77.59 | 75.71 |
|  | Agent A | 82.69 | 83.68 | 81.69 |
|  | Agent B | 49.77 | 49.77 | 49.77 |
|  | EviSearch | 77.61 | 78.31 | 76.92 |

**Table 3.** Evidence verification and modality-specific error rates across model families. Percentages are reported on the indicated denominator.

| Model | Verifier Coverage | Reproduction Among Checked | Text Error | Table Error | Figure Error |
| --- | --- | --- | --- | --- | --- |
| Qwen | 50.68 | 97.63 | 15.41 | 17.12 | 40.00 |
| Mistral | 61.73 | 82.95 | 23.26 | 37.69 | 56.38 |
| Gemma | 53.76 | 81.68 | 31.41 | 3.17 | 35.42 |

Gemma showed the opposite pattern. Correctness decreased from 89.10% for B1 to 83.76% for EviSearch, and completeness decreased from 88.16% to 83.65%. The results therefore reflect changes in both the extracted clinical value and the information required by the column definition rather than completeness alone.

Accuracy is the mean graded score across cells. Each cell is scored using correctness and completeness. Correctness measures whether the extracted answer is substantively correct. Completeness measures whether the answer contains the information required by the column definition.

### 2.5 Disagreement and review signals

Agreement between the two extraction agents was associated with higher EviSearch accuracy, while disagreement was associated with higher error rates. Figure 3 shows this separation across the four model families.

**Fig. 3.**
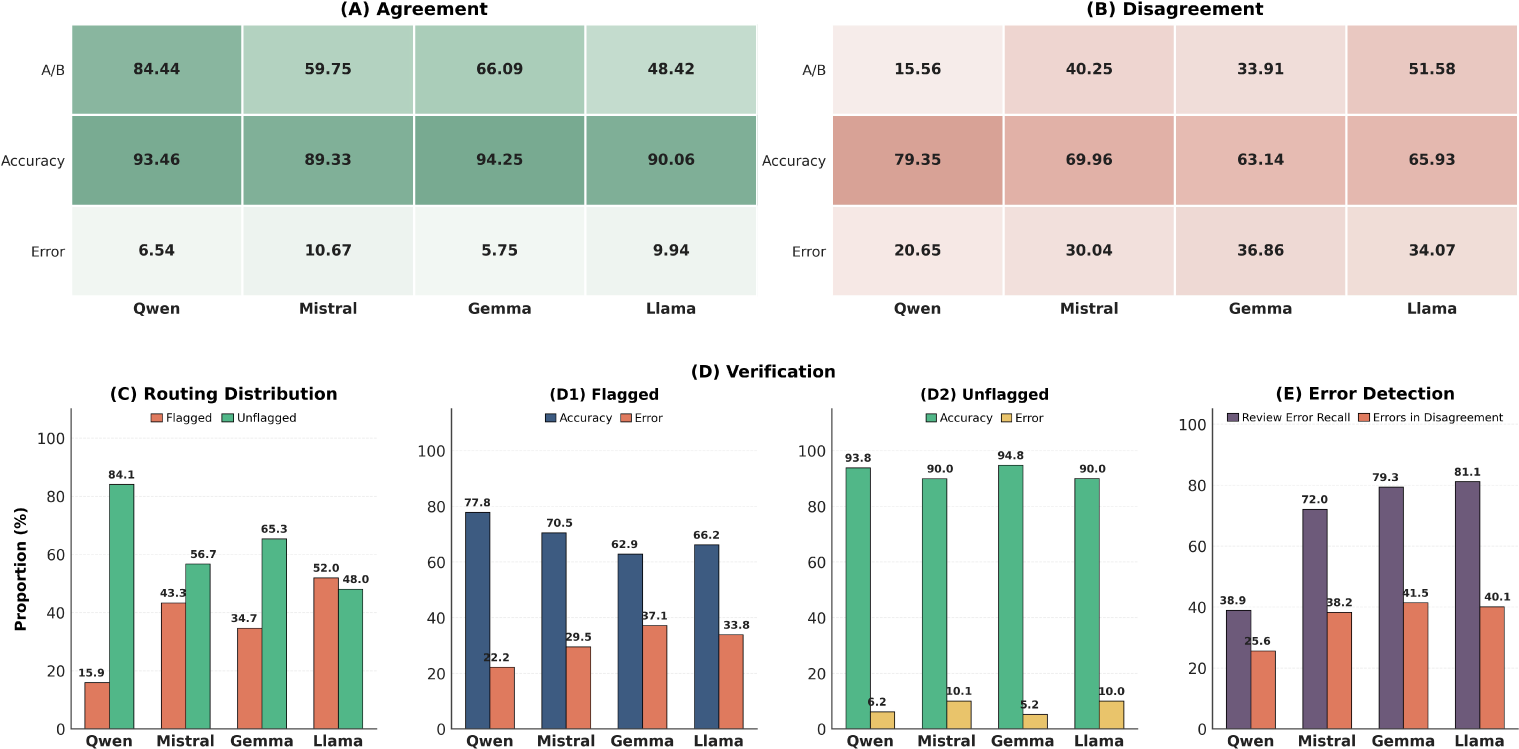
Agreement, disagreement, and review signals across model families. Panels (a) and (b) show the distribution of Agent A and Agent B agreement and disagreement and the corresponding EviSearch accuracy and error rates. Panel (c) shows the proportion of cells flagged and unflagged for review. Panel (d) shows flagged and unflagged accuracy and error rates together with review error recall.

The proportion of cells requiring review also differed across models. Qwen had 15.56% disagreement and 15.94% of cells were flagged for review. Mistral had 40.30% disagreement and 43.31% of cells were flagged. Gemma had 33.91% disagreement and 34.66% of cells were flagged. Llama had the highest disagreement at 51.58%, with 51.95% of cells flagged for review.

The review queue contained 38.93% of all EviSearch errors for Qwen with a 95% CI of 27.21 to 51.52, 72.05% for Mistral with a 95% CI of 59.84 to 81.60, 79.34% for Gemma with a 95% CI of 74.44 to 83.50, and 81.12% for Llama with a 95% CI of 70.25 to 89.75. Flagged cells were less accurate than unflagged cells across the evaluated model families. Disagreement and review status therefore identify portions of the extracted clinical evidence with a greater concentration of errors. They are used as review signals and not as guarantees of correctness.

### 2.6 Evidence provenance

Evidence provenance was evaluated separately from disagreement-based review. The attribution verifier checked 674 Qwen cells, 821 Mistral cells, and 715 Gemma cells.

Of these, 658 Qwen cells, 681 Mistral cells, and 584 Gemma cells were independently reproduced. This corresponds to verifier coverage of 50.68%, 61.73%, and 53.76%, with reproduction among checked cells of 97.63%, 82.95%, and 81.68%, respectively.

Figure 4 shows the number of cells checked by the attribution verifier and the number for which the independent verifier reproduced the extracted value across Qwen, Mistral, and Gemma.

**Fig. 4.**
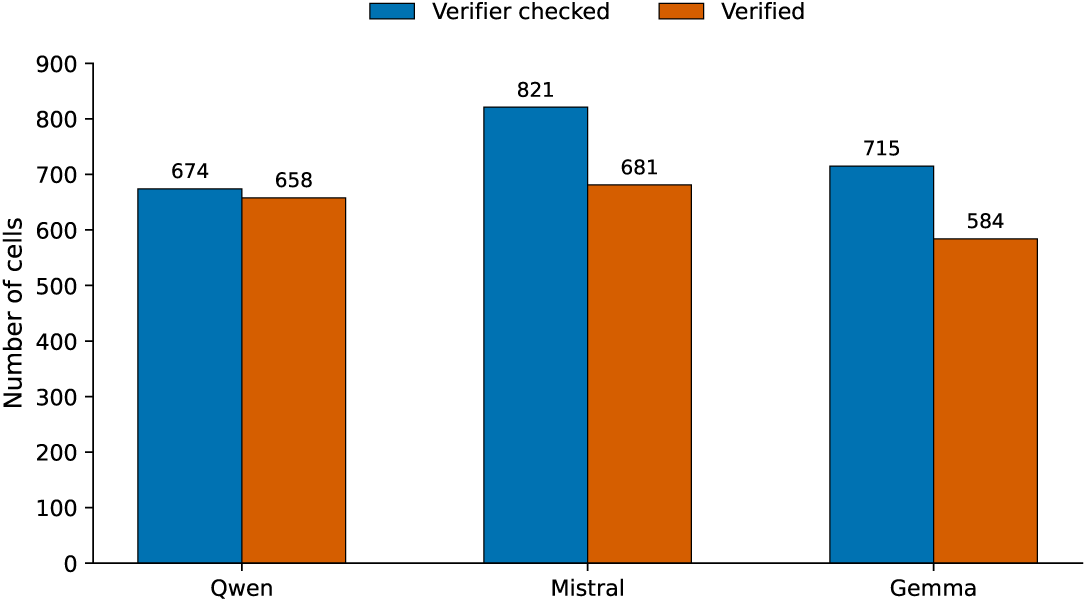
Attribution verification across model families. Bars show the number of cells checked by the attribution verifier and the number for which the independent verifier reproduced the extracted value.

The modality-specific results show differences in error rates for evidence extracted from text, tables, and figures. For Qwen, error rates were 15.41% for text, 17.12% for tables, and 40.00% for figures. For Mistral, the corresponding rates were 23.26%, 37.69%, and 56.38%. For Gemma, they were 31.41%, 3.17%, and 35.42%.

Verifier coverage is the proportion of all cells checked by the attribution verifier. Reproduction among checked cells is the proportion of checked cells for which the independent verifier reproduced the extracted value. These measures are reported separately because the verifier did not inspect every cell.

### 2.7 Cross-model reliability profile

Taken together, the cross-model results show that extraction accuracy and reliability signals do not move uniformly across model families. Qwen showed the strongest overall EviSearch performance and the lowest disagreement rate, while Mistral, Gemma, and Llama exhibited substantially higher disagreement and larger review queues. At the same time, disagreement and review flags consistently concentrated a greater share of extraction errors across all four model families.

Evidence verification showed a similar pattern. Qwen had the highest reproduction rate among verifier-checked cells, while Mistral and Gemma showed lower reproduction despite substantial verifier coverage. These results indicate that final extraction accuracy, agent agreement, review signals, and evidence reproduction provide complementary views of system reliability rather than interchangeable measures.

## 3 Discussion

This study addresses a practical problem in evidence-based medicine: how to accelerate extraction without making the evidence base harder to inspect. The results indicate that extraction quality should be considered together with source traceability and the ability to identify cells that warrant expert attention. EviSearch is therefore best understood as decision support for evidence curation, not as a replacement for clinical or methodological judgment.

The cross-model findings also show that no single computational model is uniformly reliable. A model can perform well on average while still producing errors in particular evidence types, populations, or reporting formats. Independent readings, source reproduction, and disagreement provide complementary information about where those errors may occur. These signals are useful for prioritization, but they do not establish clinical truth and should not be used to suppress review of apparently low-risk cells when the evidence is consequential.

The broader implication is that clinical evidence systems should report an audit trail alongside extraction performance. Reviewers need to know which publication passage supports a value, whether that passage was independently reproduced, and how the system handled figures, tables, and incomplete reporting. This approach could support living reviews and guideline updates by making revisions easier to verify and by preserving the reasoning behind changes to definitions or evidence values.

Several limitations temper these implications. The benchmark covers one disease area and one evidence-table schema, and many semantic comparisons rely on a blinded large language model judge rather than independent clinician adjudication of every cell. The review analysis is simulated from reference annotations and does not measure prospective reviewer workload, correction time, or inter-reviewer agreement. Some values depend on distributed or visual evidence that cannot be represented by a single highlighted span. Finally, the tested model families and deployment conditions do not establish universal model independence.

Future work should evaluate EviSearch prospectively with clinical and methodological reviewers, measuring review time, correction burden, inter-reviewer agreement, and downstream changes to systematic reviews or guidelines. External validation should span additional diseases, trial designs, languages, publishers, and evidence schemas. Further studies should also test calibration of disagreement and verifier signals, compare alternative evidence-location methods for figures and complex tables, and assess whether the system improves the timeliness and reproducibility of living evidence updates without changing clinical conclusions.

## 4 Methods

Given a publication *D* and a schema *S* = *{(c_i_, d_i_)}^n^_i=1_* containing column names and natural-language definitions, EviSearch returns either a value *v_i_*or an explicit nonreporting label. A value is accompanied by an attribution *a_i_* = (*p_i_, q_i_, m_i_*), where *p_i_*is a source page, *q_i_* is a supporting quotation, and *m_i_*is the evidence modality: text, table, or figure. The benchmark schema contains 133 columns covering trial design, treatment arms, baseline characteristics, efficacy, subgroups, and adverse events.

### 4.1 Document ingestion and extraction agents

At ingestion, a publication is parsed into page-level text with modality labels, rendered page images, and a document-specific dense retrieval index. The PDF Query Agent (A) receives the complete parsed document and, when available for the model configuration, page images, and answers a column batch in a single structured call. The Search Agent (B) begins without the full document and retrieves page-level text through dense search and page lookup tools. The two agents produce independent candidates and do not read one another’s outputs. The Reconciliation Agent first performs an independent extraction using retrieval and page-image tools, then adjudicates the three readings after the candidates are revealed.

### 4.2 Attribution verification and admission

The attribution verifier receives a candidate and its cited pages, formulates its own answer before comparing it with the candidate, and returns a page, quotation, modality, and reproduction status. A value is admitted with its attribution after this check. If the verifier does not reproduce the value, the value remains admissible with the extractor’s quotation but is marked uncorroborated and routed to review. For non-reported values, admission requires an explicit basis describing the pages examined and what they report.

### 4.3 Review and improvement with use

The review queue prioritizes cells where A and B disagree or where attribution was not independently reproduced. Reviewer corrections are stored with a rationale. Schema-level interpretations update versioned definitions or knowledge notes, while extraction notes describe how values are found and assembled. Every run records the knowledge fingerprint in force.

### 4.4 Evaluation metrics and statistical analysis

Each extracted cell was evaluated for correctness and completeness on *{*0, 0.5, 1*}*. Correctness measures whether the extracted value is substantively consistent with the reference annotation, while completeness measures whether all information required by the schema definition is present. Cell-level accuracy is defined as

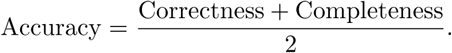

System-level accuracy is the mean cell-level accuracy across the 1,330 cells for each model condition. A binary extraction error was defined as any cell with correctness *<* 1 or completeness *<* 1.

Agent agreement was determined using the same normalized comparison procedure used in the disagreement analysis. Review error recall was defined as the proportion of all EviSearch error cells that were routed to the review queue. Verifier coverage was the proportion of all cells examined by the attribution verifier, and reproduction among checked cells was the proportion of checked cells for which the verifier independently reproduced the extracted value.

Statistical uncertainty was estimated using a publication-clustered bootstrap. For each model family, the ten publications were sampled with replacement, retaining all 133 cells from each selected publication. We generated 10,000 bootstrap replicates using seed 20260927 and report percentile 95% confidence intervals from the 2.5th and 97.5th percentiles. Differences between EviSearch and B1 were evaluated using paired bootstrap samples in which the same resampled publications were used for both systems.

### 4.5 Research questions and benchmark

The primary question is whether attribution, disagreement, verification, and selective review remain informative across tested model families. The benchmark comprises ten randomized-trial publications in metastatic hormone-sensitive prostate cancer, each represented by the same locked 133-column schema and a clinician-annotated reference row. Correctness and completeness are scored on *{*0, 0.5, 1*}*, and accuracy is the mean of the two scores over all cells. The semantic scoring component uses a blinded large language model (LLM) judge under a written rubric; identical predictions are cached and judged once.

### 4.6 Trust signals and review

For each cell, we record A/B agreement, verifier reproduction, final correctness, citation status, and review status. We report disagreement frequency, review-flag frequency, review error recall, verifier coverage, reproduction among checked cells, and modality-specific error rates. Review error recall quantifies the proportion of binary EviSearch errors routed to review and does not represent measured clinician workload. Statistical uncertainty for review error recall was estimated using the publication-clustered bootstrap described above.

### 4.7 Reproducibility and software verification

The released implementation documents the extraction pipeline, model configurations, retrieval settings, prompt definitions, evaluation procedure, and statistical analysis used in this study. The repository includes the locked 133-column mHSPC schema, frozen extraction rules, benchmark data, stage-wise prompts, evaluation rubric, and analysis scripts. Exact model identifiers are provided for all evaluated model families, together with the shared Qwen3-Embedding-8B retrieval model and top-*k* = 5 retrieval setting. The same scoring rubric and evaluation procedure were applied across all model families.

## Data and code availability

The benchmark data, reference annotations, schema, EviSearch source code, extraction prompts, frozen extraction rules, evaluation rubric, experiment scripts, and statistical analysis code used in this study are available in the anonymous project repository (https://anonymous.4open.science/r/Reliable-extract ion-of-clinical-trial-evidence-D7F8/).

## Supplementary Information

### Prompt Templates

For readability, we summarize the fixed stage-wise prompts used by EviSearch here. The released code contains the complete prompt strings, structured output schemas, tool definitions, and frozen extraction rules used in the experiments. Dynamic document text, column definitions, candidate answers, and evidence are represented by placeholders.

### Single-Pass Baseline

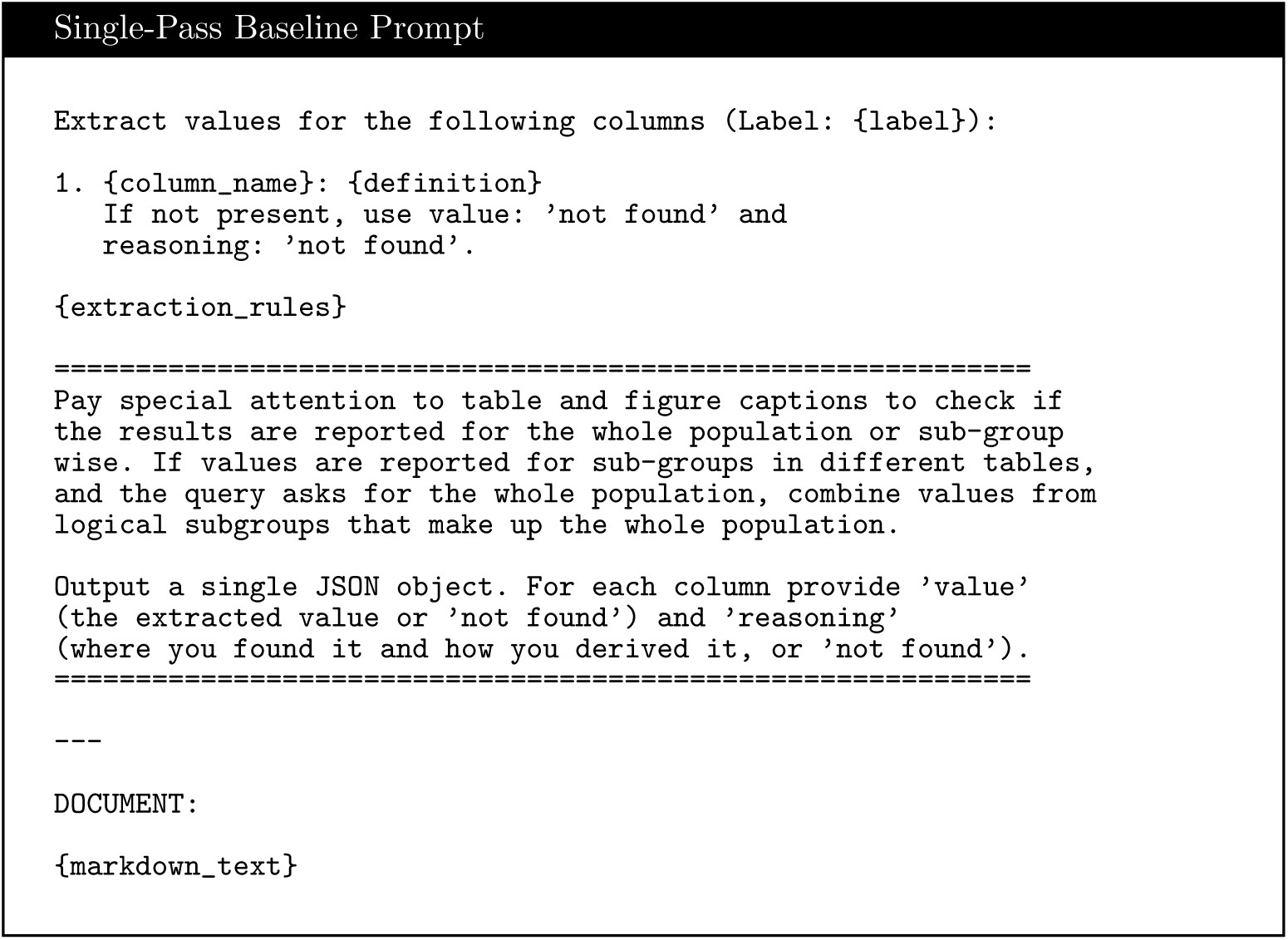

#### Baseline Extraction

The baseline receives the parsed publication and requested schema fields in a single model call. A structured JSON schema constrains the response to contain a value and reasoning for every requested column.

#### PDF Query Agent

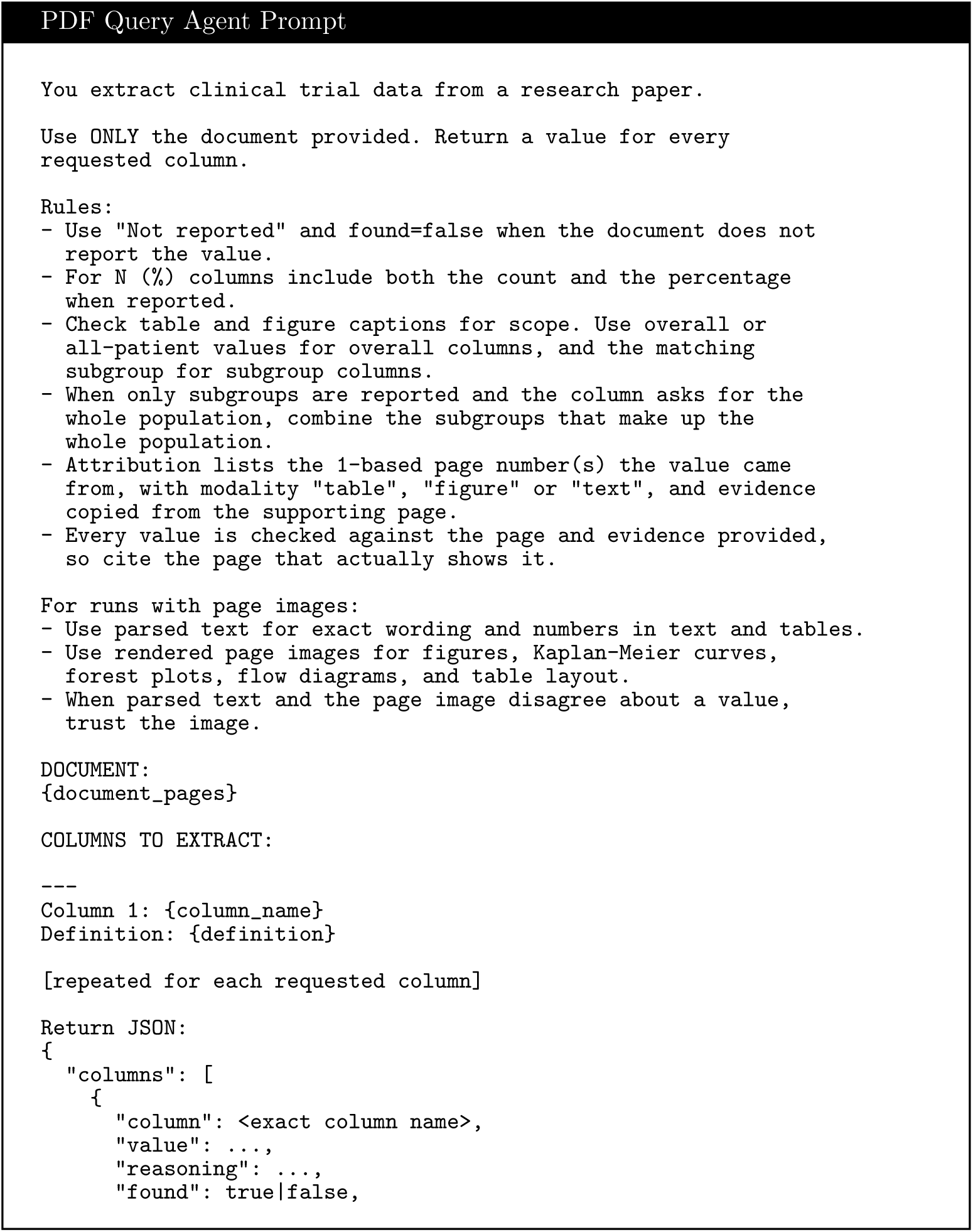
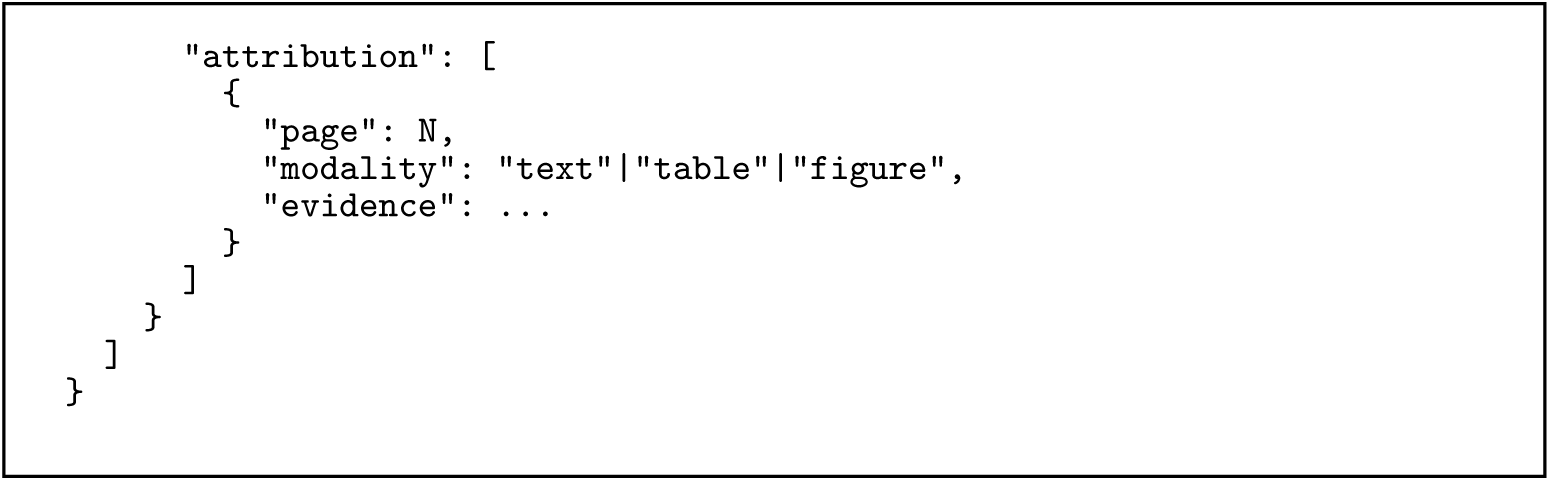

#### Document Input

The PDF Query Agent receives the complete parsed publication. Qwen and Mistral additionally receive rendered page images, while Gemma and Llama use parsed text only. The requested columns are processed in batches and the response is constrained by a structured output schema.

#### Search Agent

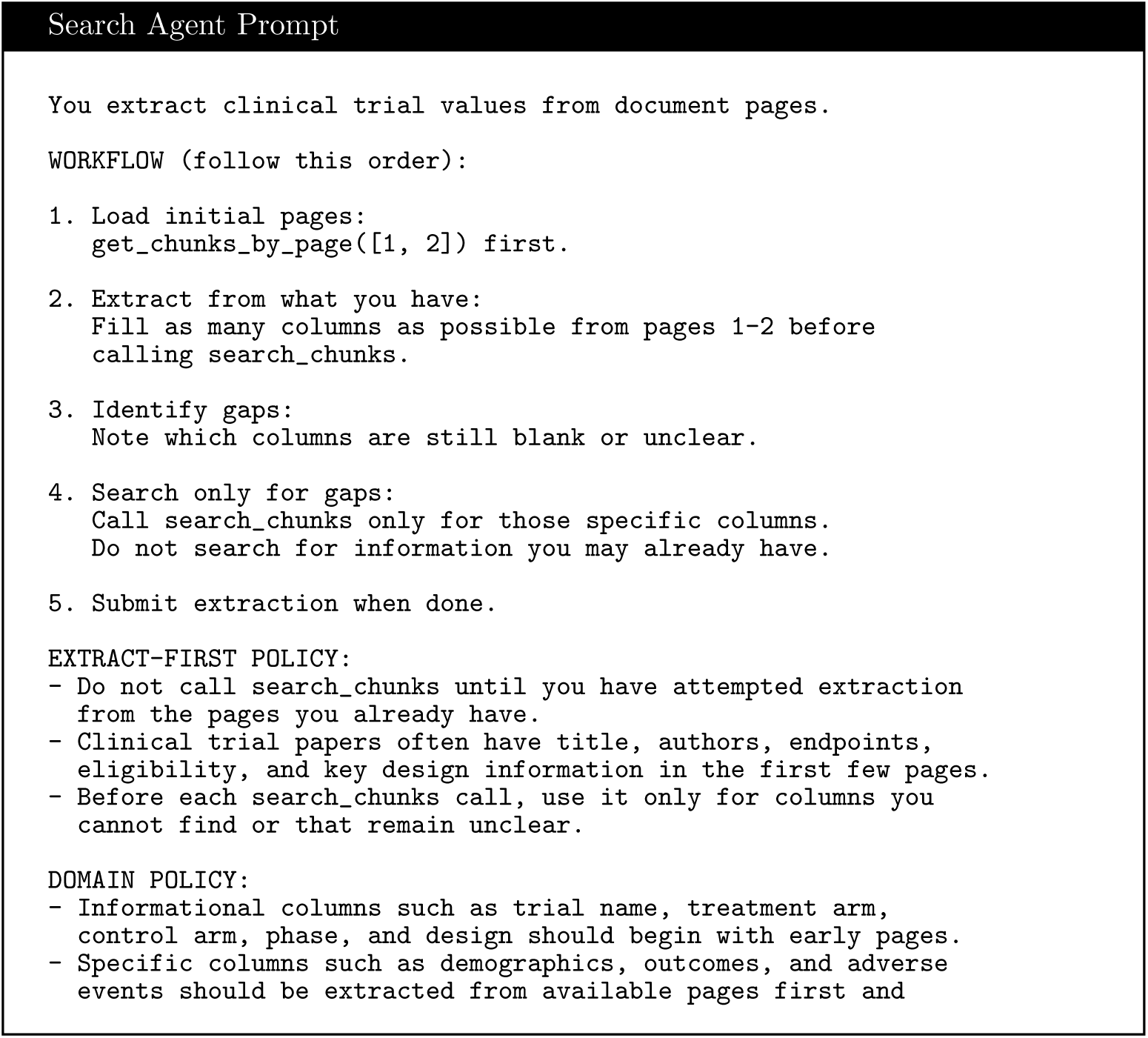
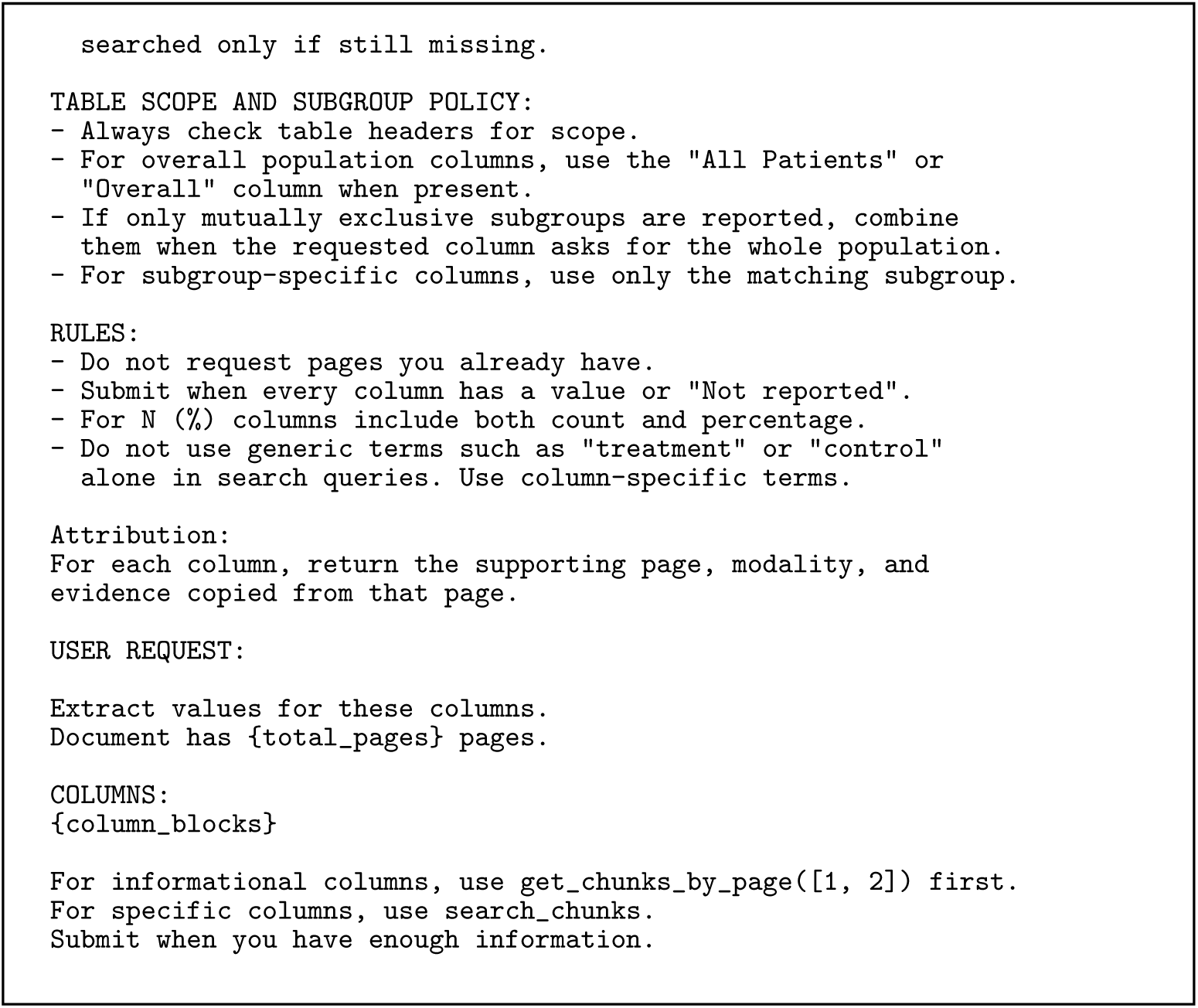

#### Search Tools

The Search Agent operates with three tools, search chunks, get chunks by page, and submit extraction. Search results contain page-level text and the agent receives the retrieved evidence before continuing extraction.

#### Reconciliation Agent

Reconciliation is performed in two phases. The first phase forms an independent reading without access to Agent A or Agent B. The second phase reveals the independent reading together with the two candidate extractions and produces the final value.

#### Independent Reading

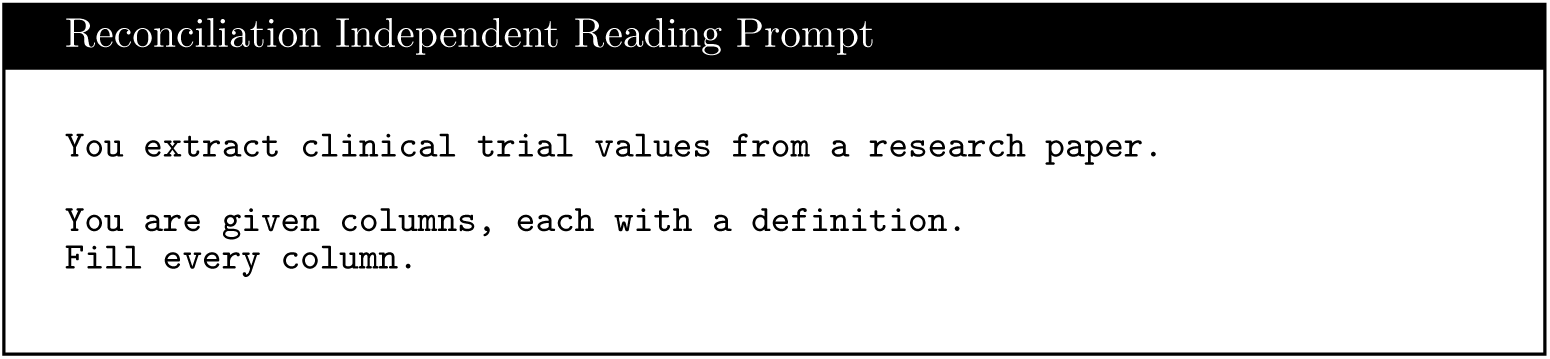
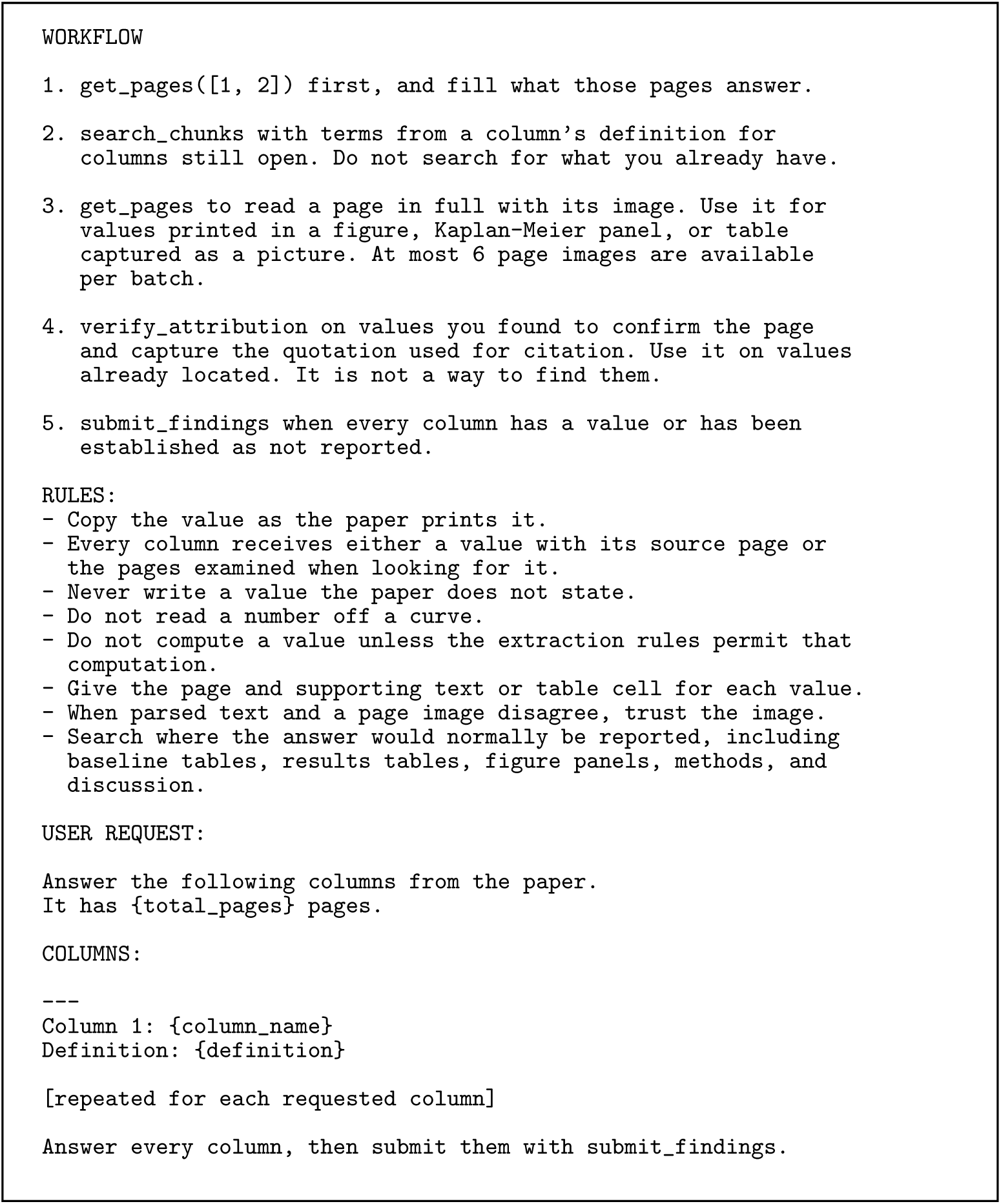

#### Final Adjudication

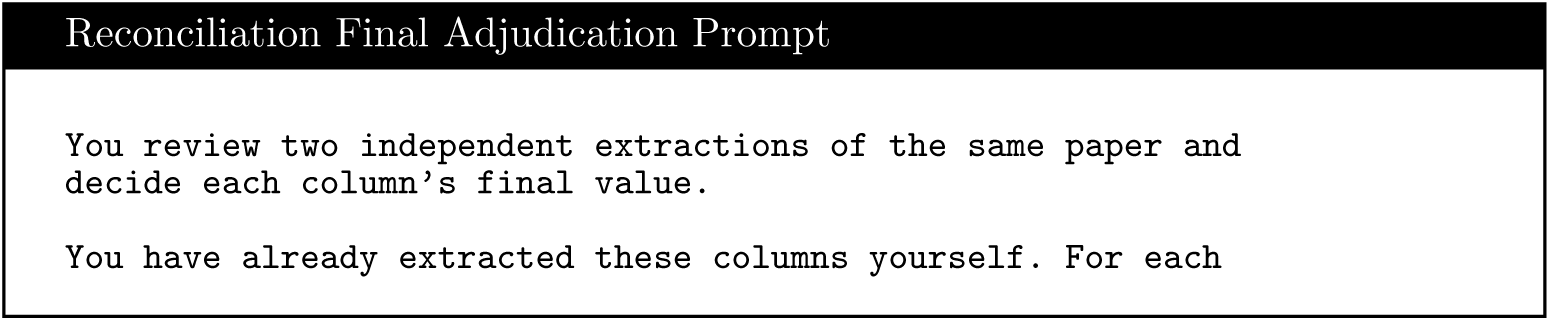
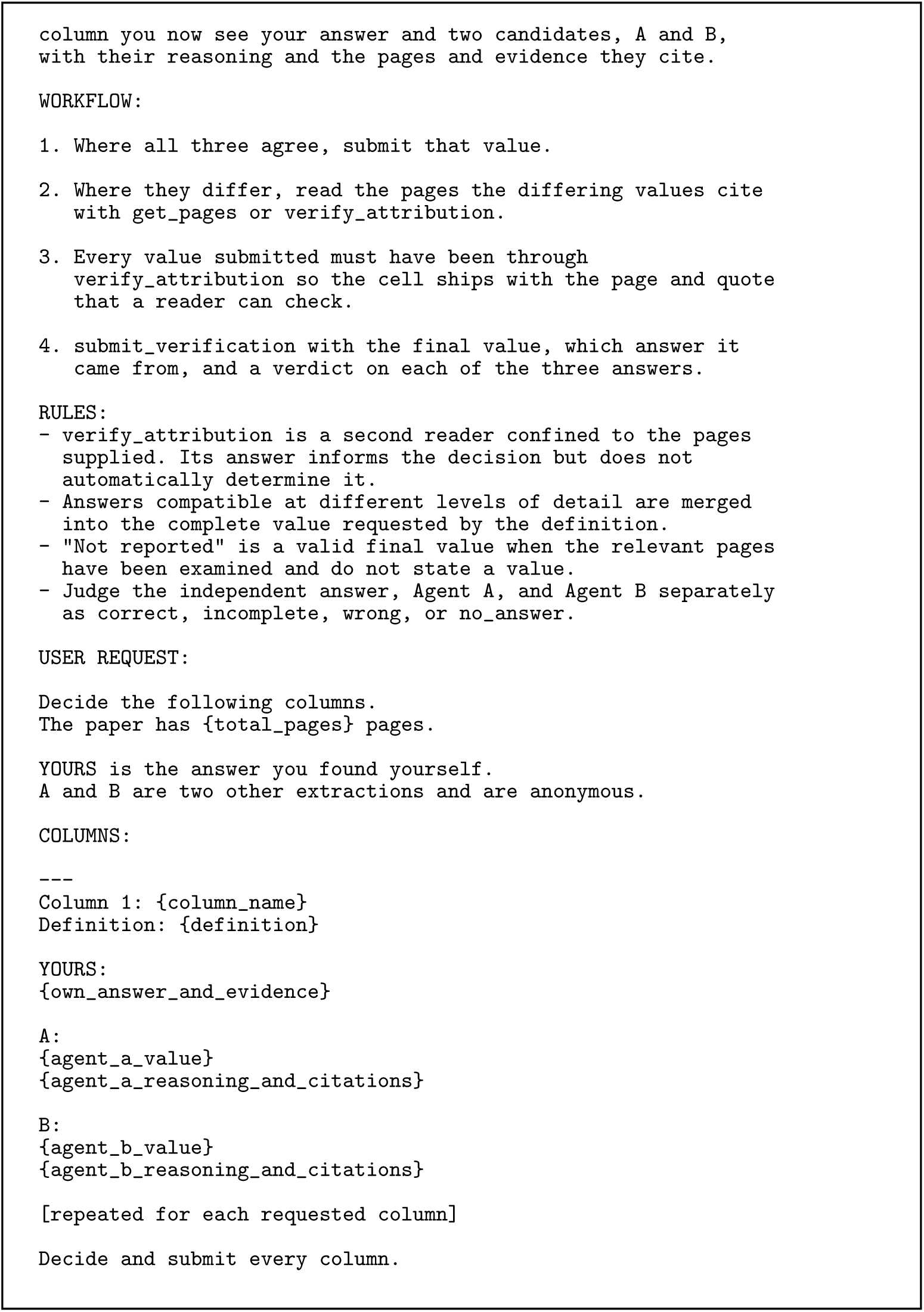

#### Reconciliation Tools

The reconciliation stage uses document search, page loading, attribution verification, and structured submission tools. Independent extraction is performed before the candidate answers are revealed.

#### Attribution Verification

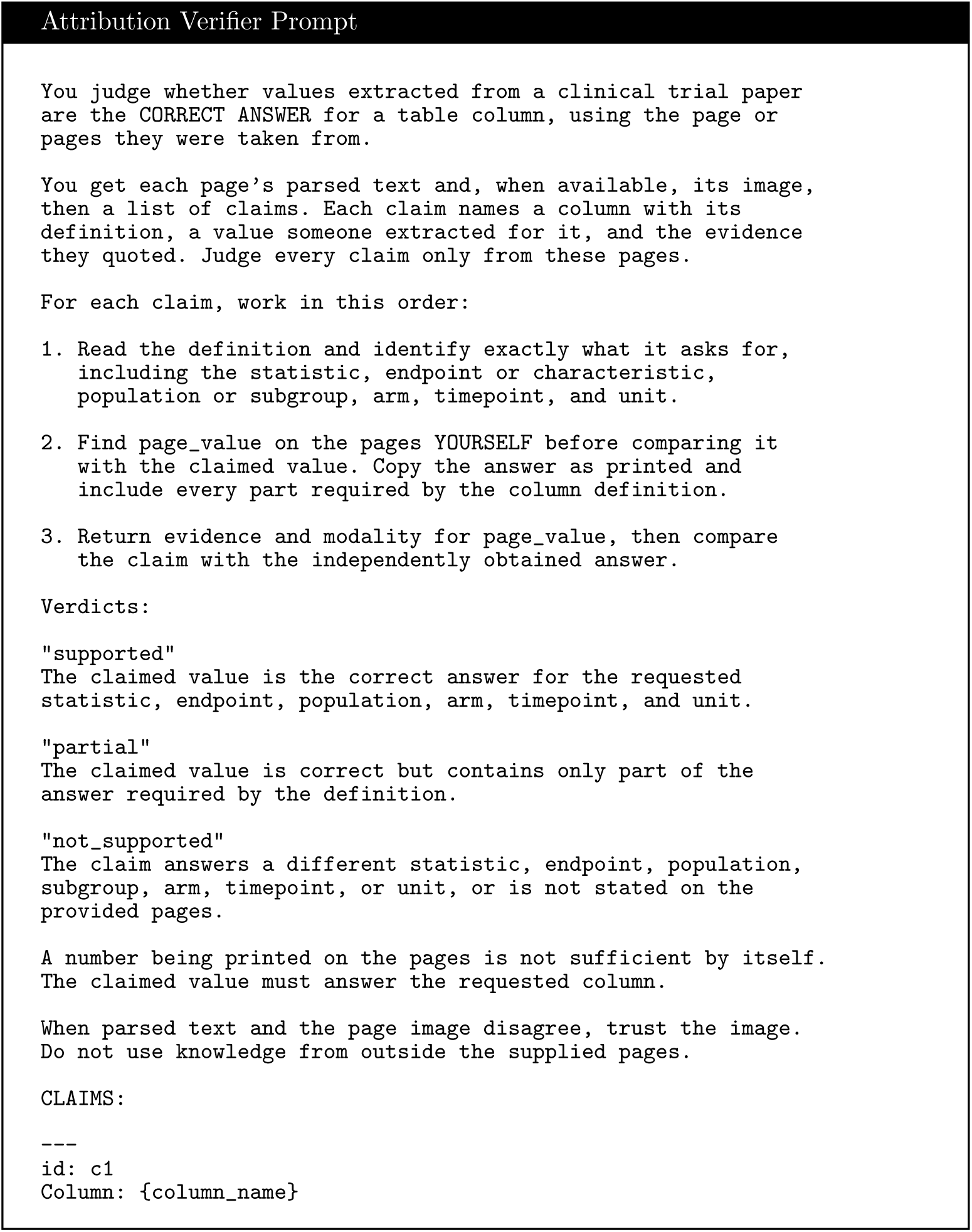
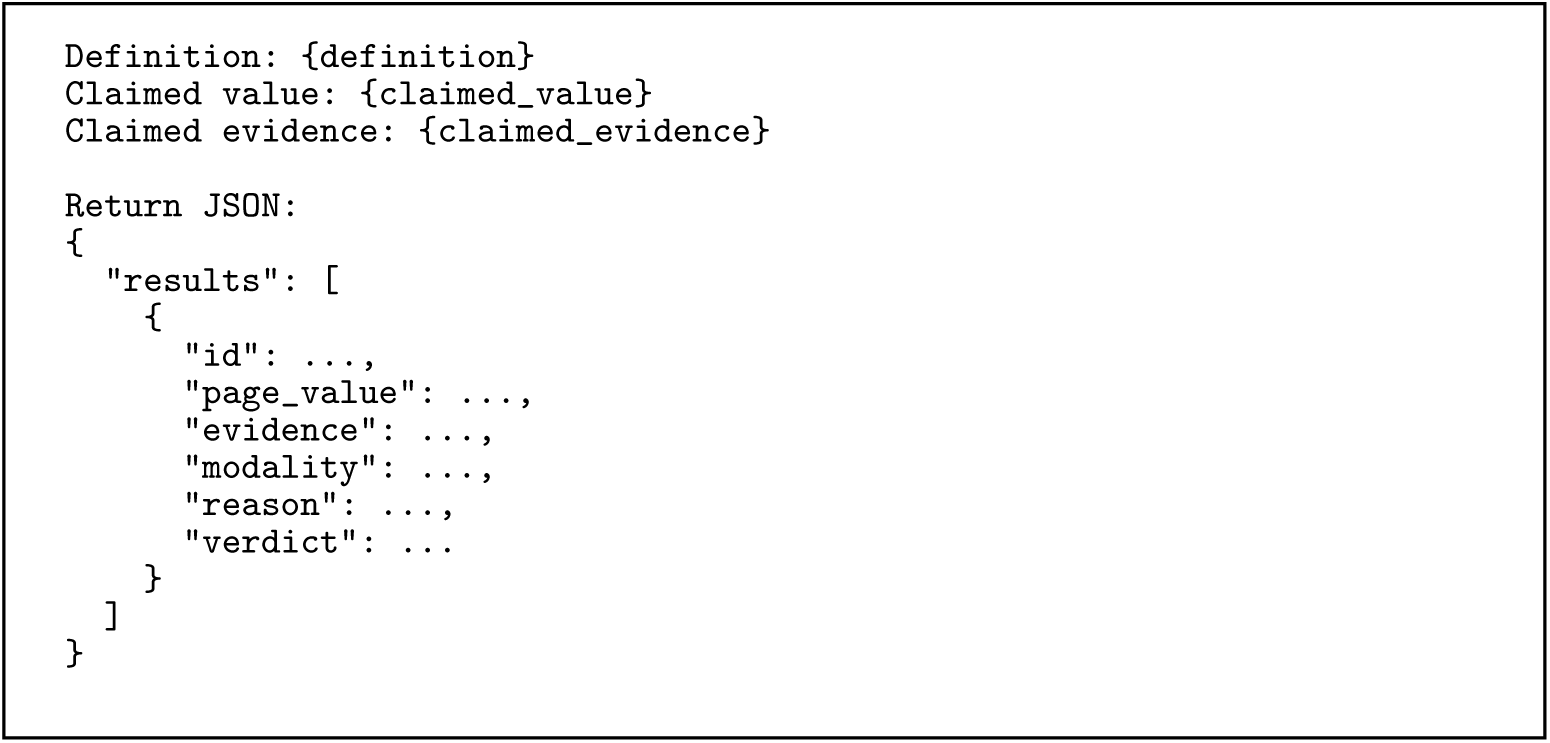

#### Verifier Output

The verifier returns one of three verdicts, supported, partial, or not supported. The verifier first generates its own answer from the cited page before comparing that answer with the candidate value.

#### Evaluation Judge

The evaluation judge receives blinded ground-truth and predicted values. It does not receive the identity of the extraction system that produced each prediction. Correctness and completeness are each restricted to *{*0, 0.5, 1*}*.

#### Exact-Match Evaluation

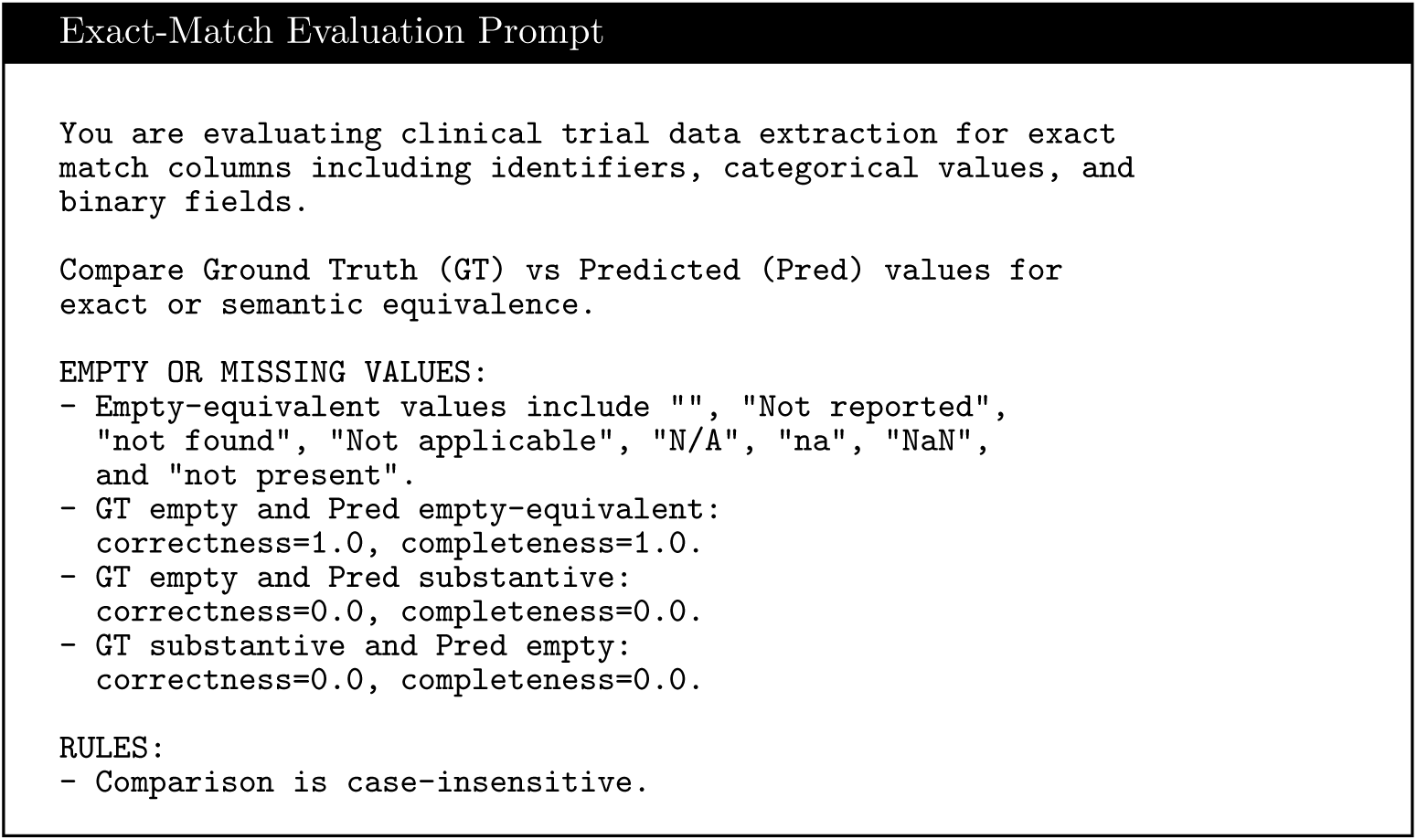
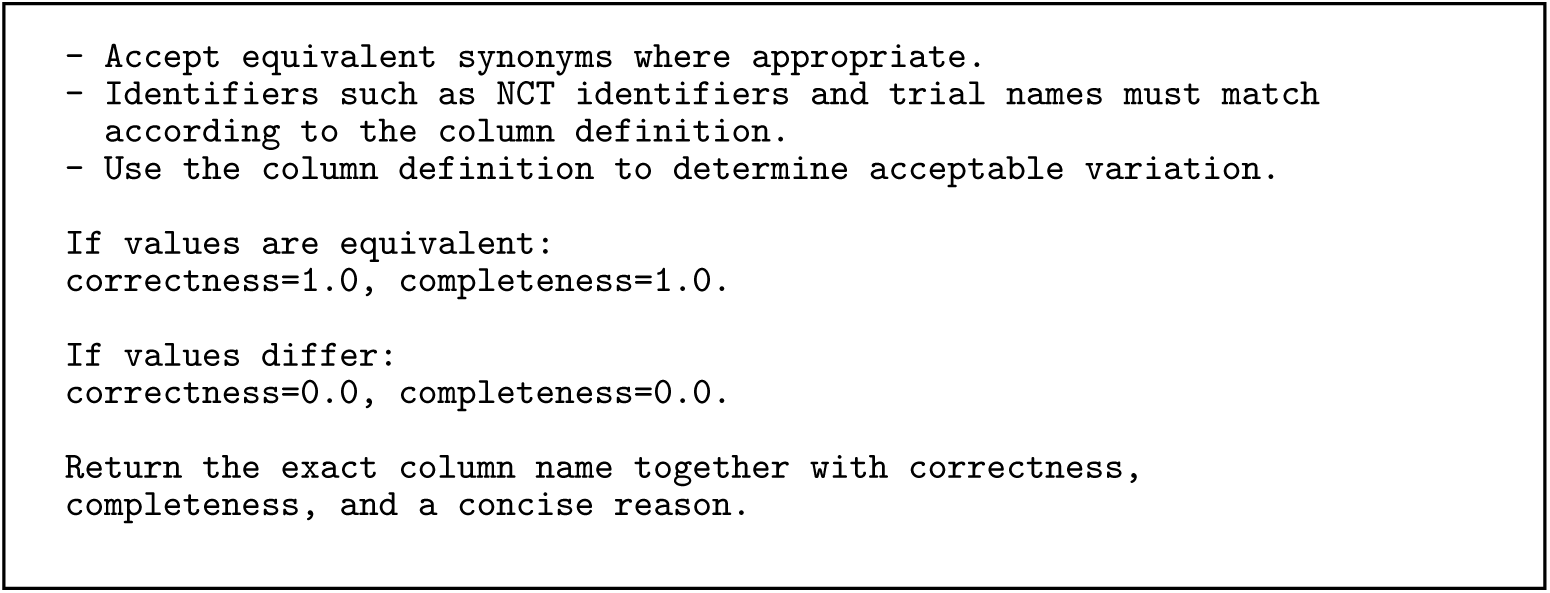

#### Numeric Evaluation

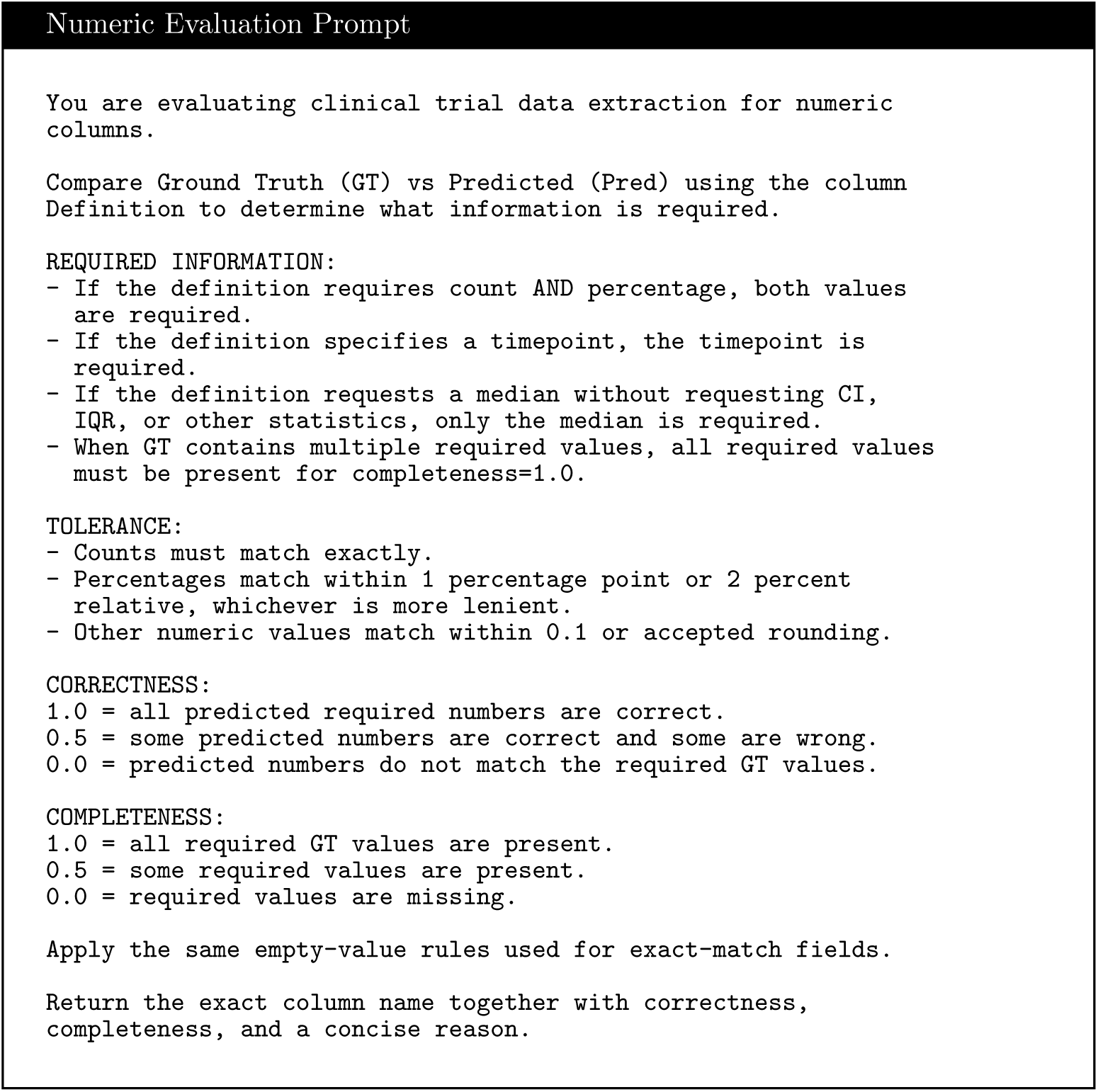

#### Structured-Text Evaluation

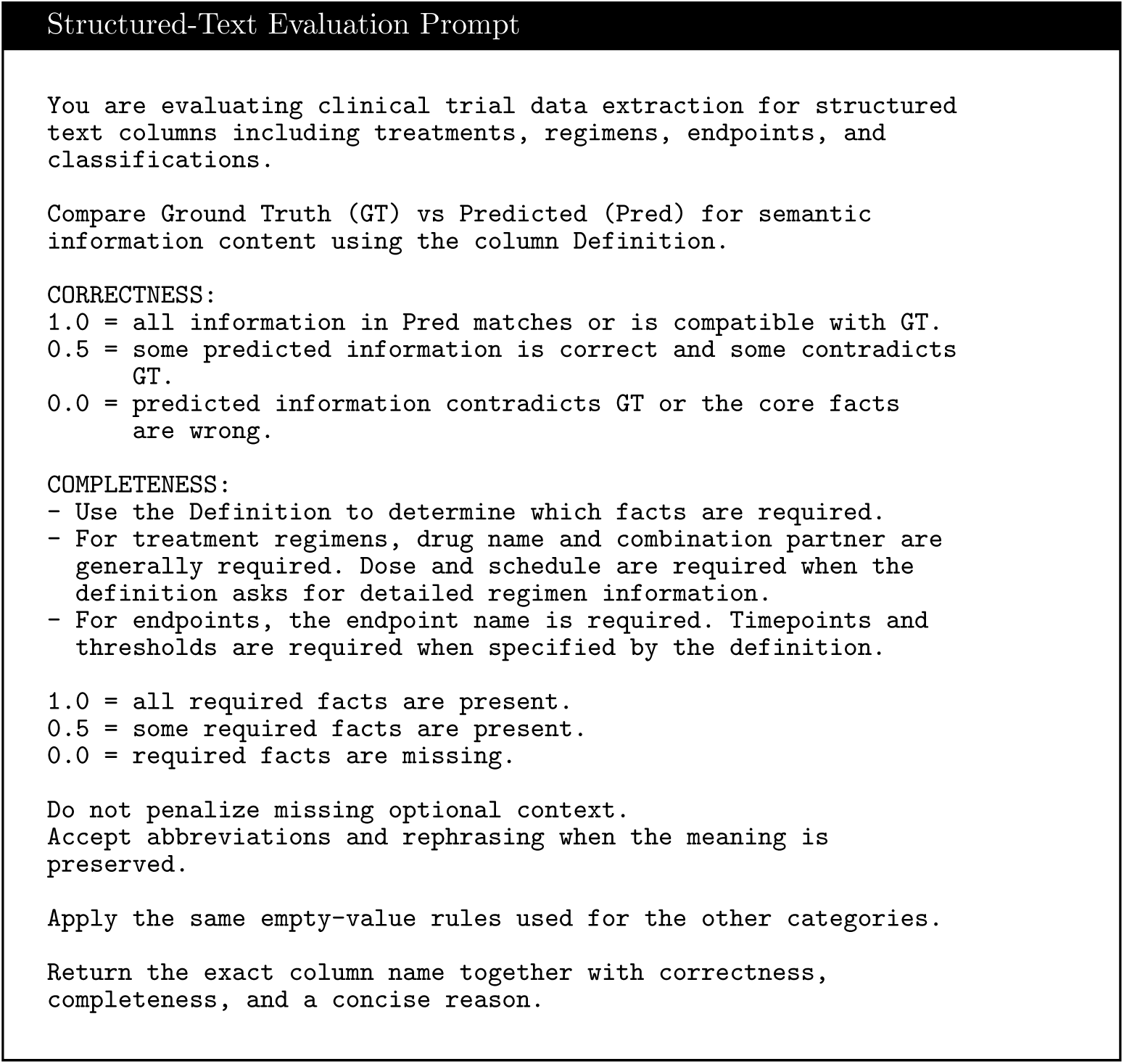

#### Scoring Output

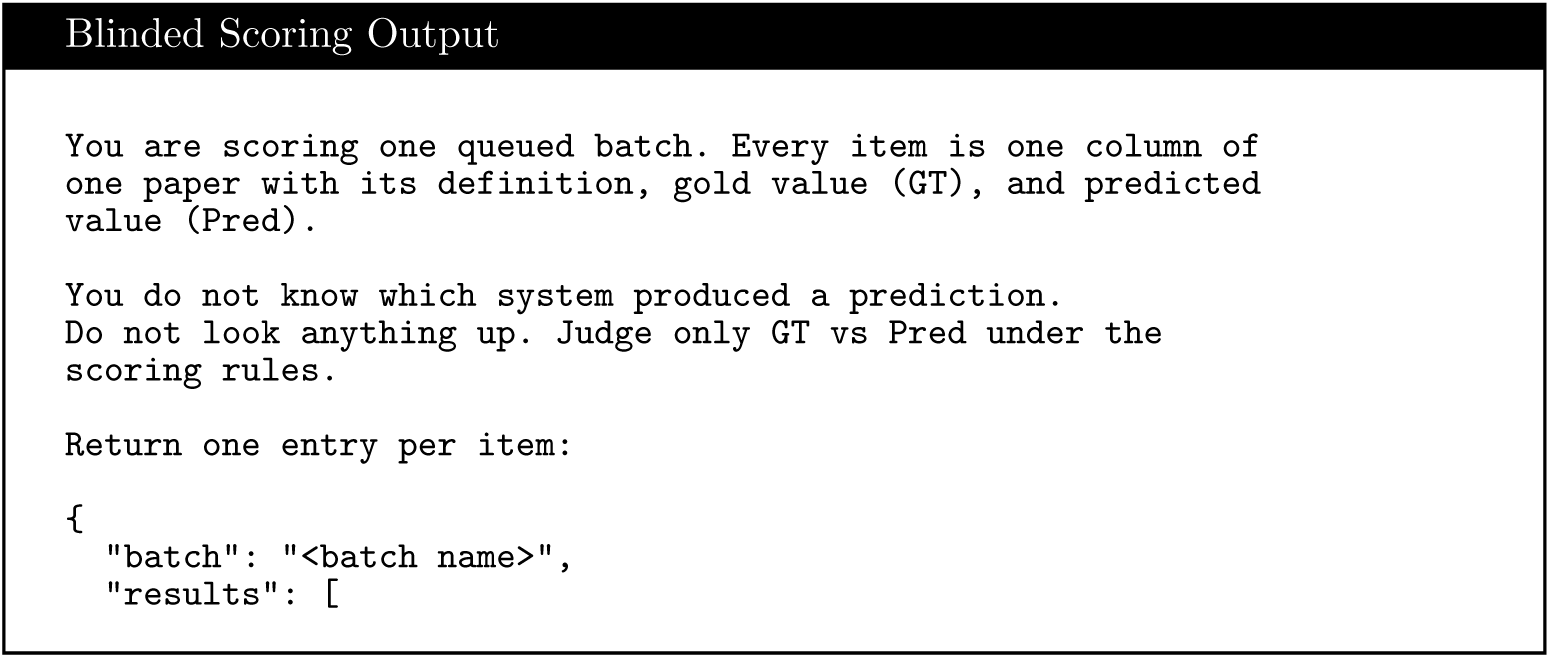
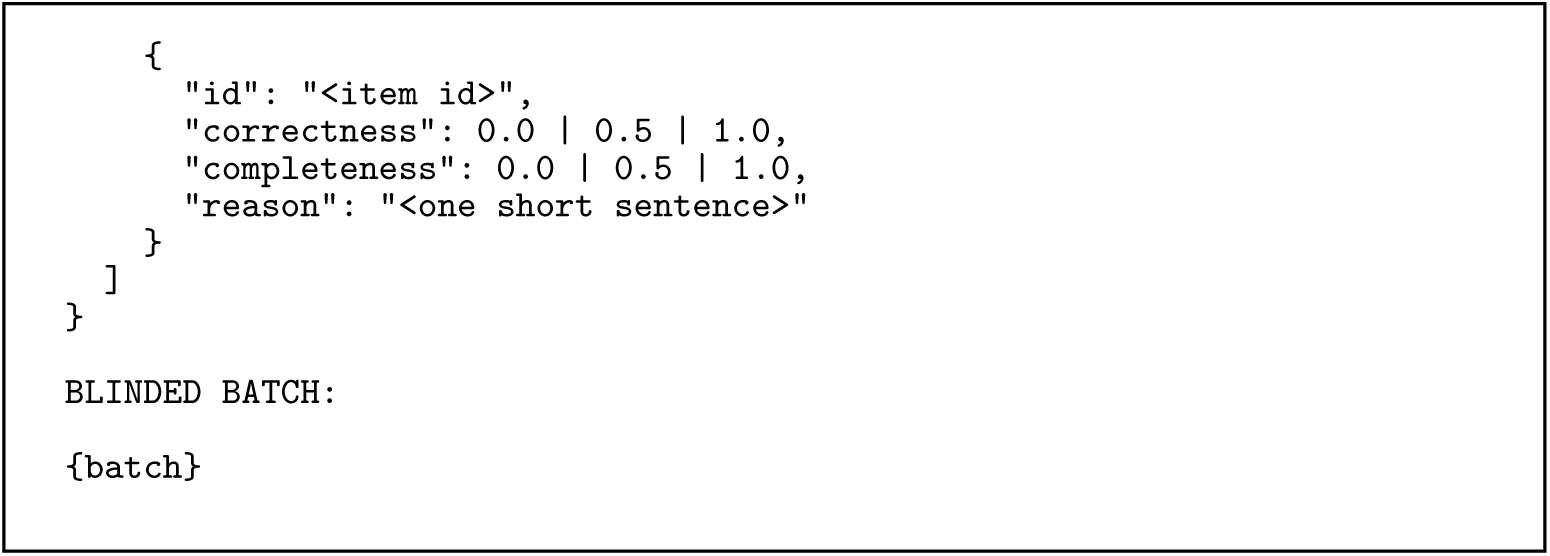

#### Evaluation Procedure

The same blinded evaluation rubric was used for all four model families. The fixed judge was Gemini 3.5 Flash with temperature 0.0. Correctness and completeness were restricted to *{*0, 0.5, 1*}*, and the scoring procedure included fixed rules for empty values, numeric tolerance, structured text, and required information.

#### Frozen Extraction Rules

The extraction stages additionally receive the frozen task-specific knowledge snapshot used in the completed runs. These rules specify consistent interpretations for clinical endpoints, population scope, treatment arms, subgroup values, and other schema-specific cases. Global rules are supplied to all relevant columns, while column-specific rules are included only for the fields to which they apply. The complete frozen rules are released with the implementation.

